# Norditerpene natural products from subterranean fungi with anti-parasitic activity

**DOI:** 10.1101/2025.01.02.631097

**Authors:** Alexandra Kolas, Yudi Rusman, Ana C.G. Maia, Jessica Williams, Fernanda G. Fumuso, Alexis Cotto-Rosario, Chidiebere Onoh, Hanen Baggar, Mary L. Piaskowski, Christian Baigorria, Raphaella Paes, Debopam Chakrabarti, Lyssa Weible, Kayode K. Ojo, Roberta M. O’Connor, Christine E. Salomon

**Author notes:** joint corresponding authors.

## Abstract

*Cryptosporidium* is a common, waterborne gastrointestinal parasite that causes diarrheal disease worldwide. Currently there are no effective therapeutics to treat cryptosporidiosis in at-risk populations. Since natural products are a known source of anti-parasitic compounds, we screened a library of extracts and pure natural product compounds isolated from bacteria and fungi collected from subterranean environments for activity against *Cryptosporidium parvum*. Eight structurally related norditerpene lactones isolated from the fungus *Oidiodendron truncatum* collected from the Soudan Iron mine in Minnesota showed potent activity and were further tested to identify the most active compounds. The availability of a diverse suite of natural structural analogs with varying activities allowed us to determine some structure activity relationships for both anti-parasitic activity as well as cytotoxicity. The two most potent compounds, oidiolactones A and B, had EC_50_s against intracellular *Cryptosporidium parvum* of 530 and 240 nM respectively without cytotoxicity to confluent HCT-8 host cells. Both compounds also inhibited the related parasite *Toxoplasma gondii*. Oidiolactone A was active against asexual, but not sexual, stages of *C. parvum*, and killed 80% of the parasites within 8 hours of treatment. This compound reduced *C. parvum* infection by 70% in IFNγ−/− mice, with no signs of toxicity. The high potency, low cytotoxicity, and *in vivo* activity combined with high production, easy isolation from fungi, and synthetic accessibility make oidiolactones A and B attractive scaffolds for the development of new anti-*Cryptosporidium* therapeutics.

## Introduction

*Cryptosporidium parvum* is a zoonotic, waterborne gastrointestinal parasite that causes significant morbidity and mortality in humans and livestock in both developing and industrialized countries [1,2]. The parasite is acquired by ingestion of food or water contaminated with oocysts, the environmentally resistant stage of the parasite [3]. *Cryptosporidium* replicates in intestinal epithelial cells, first undergoing asexual replication followed by sexual development [4] and production of oocysts that are subsequently released in the feces [3]. The infection is perpetuated by autoinfection with thin-walled oocysts; the diarrhea caused by *Cryptosporidium* can last up to 2-3 weeks, resulting in significant morbidity, and sometimes mortality [5,6].

In the United States, *Cryptosporidium* was responsible for the largest outbreak of waterborne diarrheal disease [7] and worldwide remains a common cause of waterborne diarrheal illness [1,8]. Currently, the chance of exposure to the pathogen is high even in industrialized countries [1,8], and is increasing [9]. Immunocompromised patients are highly susceptible to *Cryptosporidium* infection and, unless the immune system can be reconstituted, the infection results in prolonged, chronic and sometimes fatal diarrhea [9]. In developing countries, *Cryptosporidium* parasites are the second most common cause of diarrhea in infants, and the only gastrointestinal pathogen associated with death in toddlers [10]. Currently, there is no effective therapeutic to treat these at-risk populations [9]. The only FDA approved drug, nitazoxinide, is ineffective in immunocompromised patients [11], and clofazimine, a repurposed leprosy drug with activity against *Cryptosporidium*, failed in clinical trials in HIV-*Cryptosporidium* co-infected adults [12]. Critically, there are few anti-*Cryptosporidium* drugs in the pipeline [13]. Given the high failure rate of candidate therapeutics as they progress through pre-clinical and clinical development, more anti-*Cryptosporidium* compounds need to be discovered and investigated [14].

*Cryptosporidium parvum* is also a significant problem in veterinary medicine. The parasite is endemic in cattle operations worldwide [15,16] and is responsible for ∼40% of neonatal calf enteritis [17], causing a significant economic impact on farmers. Moreover, cattle and dairy farms serve as an environmental reservoir of this zoonotic pathogen; contamination of recreational waters by farm runoff has been documented [13]. Currently, there are no drugs available to treat cryptosporidiosis in neonatal ruminants [17]. Thus, targeted development of effective therapeutics for cryptosporidiosis remains both a veterinary and medical imperative.

Throughout history, natural products have played a pivotal role in the discovery and development of treatments for parasitic disease. Plants, mushrooms, and plant-based remedies have been used by traditional healers as essential medicines to treat infectious diseases [18]. The discovery and development of quinine from the bark of cinchona “fever trees” [19] and artemisinin from sweet wormwood (*Artemesia annua*) [20] revolutionized treatments for malaria across the globe. The more recent example of the discovery of the avermectins from *Streptomyces avermitilis* [21] and subsequent development into ivermectin for the treatment of nematode infections, highlights the potential of microbial sources of bioactive lead compounds for drug development [22]. The rich diversity of both chemical structures and source organisms has proven to be a valuable resource for the discovery of potent anti-parasitic compounds with novel mechanisms of action [23,24].

As part of a program to identify new natural product inhibitors of *C. parvum*, we screened a library of 192 crude extracts, enriched fractions and pure compounds from fungi and bacteria collected from a subterranean iron mine in Tower, Minnesota. The source microbial strains were selected for this pilot screen due to their anti-fungal activity identified in a previous biocontrol study [25]. We found several related norditerpene compounds possessing anti-*Cryptosporidium* activity; two of these compounds exhibited submicromolar activity against this parasite and the related apicomplexan, *Toxoplasma gondii*. Atypically for most natural products, these two active oidiolactones are produced in high titers by the *O. truncatum* fungus (>60 mg/Kg) and are the most abundant secondary metabolites produced in solid culture by this strain under our conditions. The plentiful availability of these compounds and potent activities encouraged us to further assess their potential via measurement of pharmacokinetic parameters and additional characterization of their anti-parasitic activity in vitro and in vivo.

## Results

### Two norditerpene lactones have potent activity against *Cryptosporidium parvum* and *Toxoplasma gondii*

A small library of 192 extracts, enriched fractions and pure compounds from 23 fungi and one bacterial species isolated from the Soudan Iron Mine, in Tower, Minnesota, was screened in an initial pilot assay against nanoluciferase expressing *Cryptosporidium parvum* [26] (Cp-NLuc)-infected HCT-8 cells at an initial dose of 5 µg/mL. A series of seven structurally related norditerpenes isolated from the fungus *Oidiodendron truncatum* showed 80-96% parasite inhibition while seven additional congeners were inactive (Fig. 1). The active compounds were retested at several concentrations to compare potencies (Fig. 1, table). The two most active compounds were oidiolactones A (**1**) and B (**7**).

**Figure 1:**
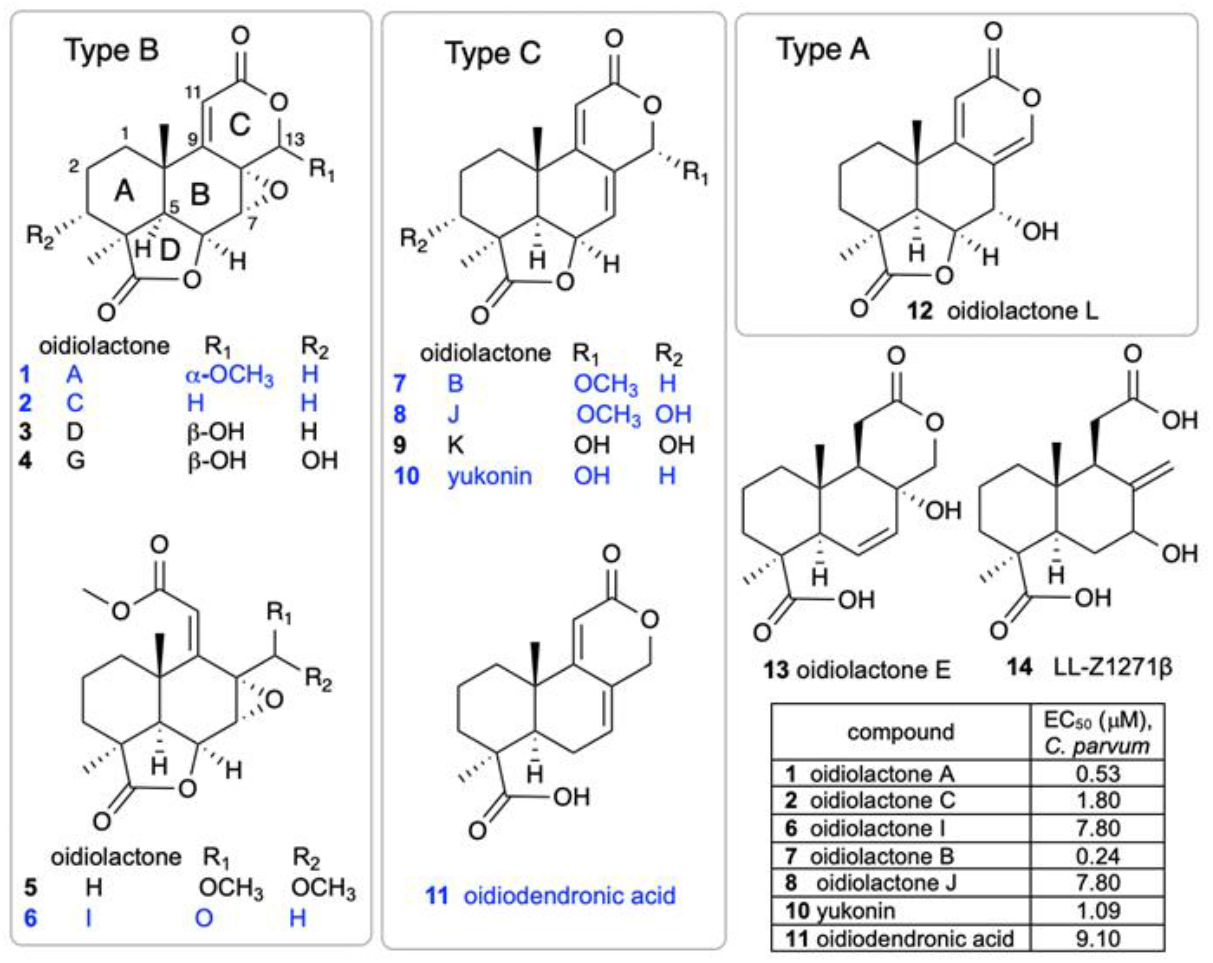
Structures of oidiolactone norditerpenes (1-14) tested against intracellular, nanoluciferase expressing-C. parvum in vitro. Active compounds are indicated in blue, and EC_50_s (µM) are shown in the table.

### Oidiolactones A and B exhibit pan anti-apicomplexan activity without toxicity to host cells

Oidiolactones A and B were tested against HCT-8 infected Cp-NLuc using an expanded range of concentrations to calculate an accurate EC_50_ (Fig. 2A). These select compounds were also tested against fibroblasts infected with the related apicomplexan parasite *Toxoplasma gondii* (luciferase and green fluorescence protein expressing RH strain *T. gondii*; TgRH-Luc:GFP; Fig. 2C). The EC_50_s of oidiolactones A and B against intracellular Cp-NLuc and TgRH-Luc:GFP were in the submicromolar range, with oidiolactone B twice as potent as oidiolactone A against both parasites (Fig. 2E). Neither compound showed any toxicity to host cells up to 100 uM when tested under the conditions of the infection assay (Fig. 2, B and D).

**Figure 2:**
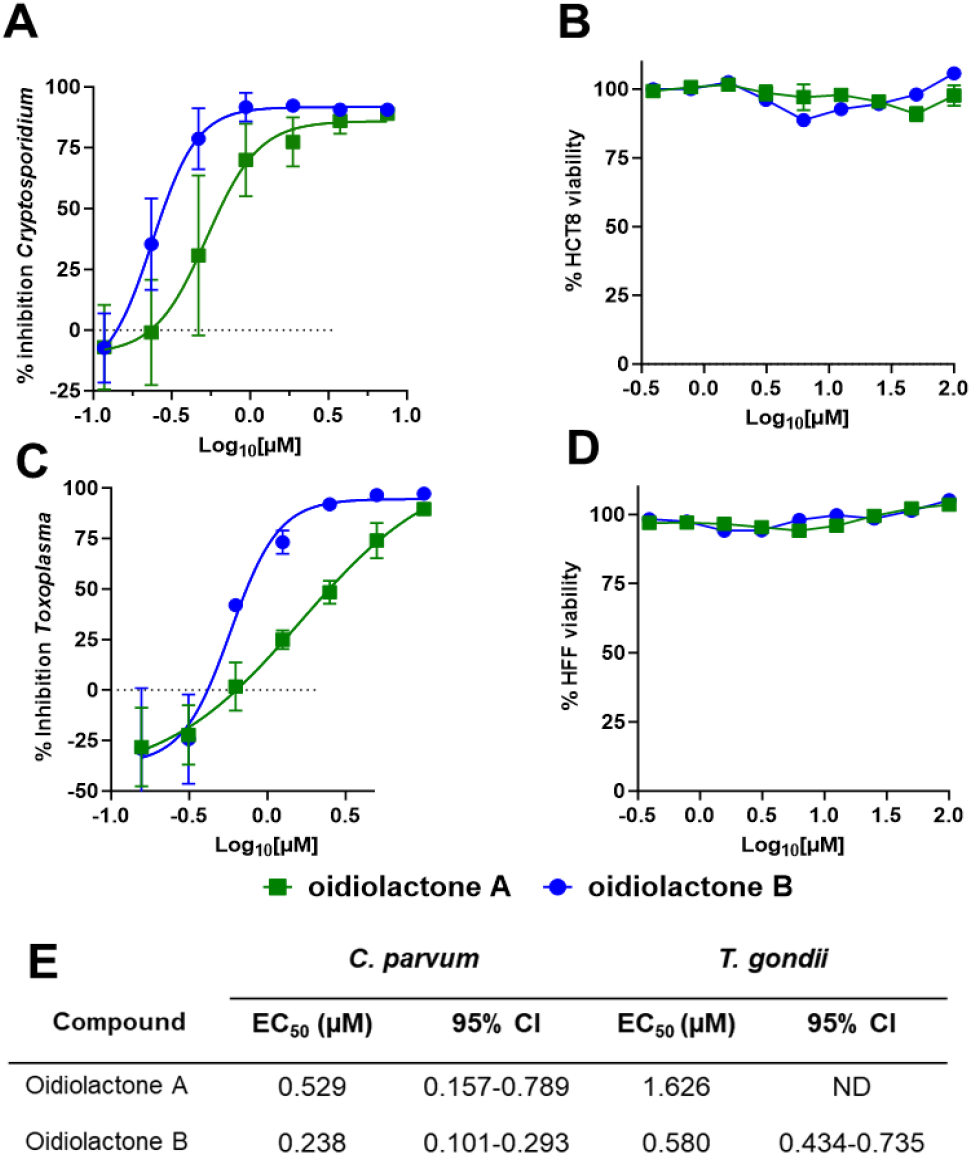
The oidiolactones have specific nM level activity against *Cryptosporidium parvum* and *Toxoplasma gondii*. Compounds were tested at concentrations ranging from 7.5 uM to 117 nM against Cp-NLuc infected HCT-8 cells (A) or 10 uM to 156 nM against TgRH-Luc:GFP infected HFF cells (C). Results are from 3 biological replicates. Confluent HCT-8 (B) and HFF (D) cells were incubated with compounds (100 uM to 390 nM) for 72 hours or 48 hours respectively. Cell viability was determined by quantification of cellular ATP levels. Results are from 2 biological replicates (E) Summary of EC_50_s and confidence intervals. EC_50_s were determined in Graphpad Prism using the log (inhibitor) vs. response–variable slope (four parameters). Error bars show the standard deviation around each point.

Since the compounds exhibited activity against two divergent apicomplexan parasites, the seven bioactive oidiolactones (Fig. 1) were also tested for activity against *Plasmodium falciparum*, the causative agent of malaria. Only oidiolactones A and B had anti-*P. falciparum* activity but at low micromolar concentrations (Suppl. Table 1). We further tested oidiolactones A and B against *Giardia lamblia* and found that they had no activity (up to 29 uM) against this protozoan.

### Oidiolactone B exhibits some cytotoxicity towards sub-confluent HCT-8 cells

To see if the compounds had any effects on actively growing cells, we tested oidiolactones A and B against sub-confluent host cells (∼40% confluency). Both oidiolactones A and B exhibited some cytotoxicity against subconfluent HCT-8 cells (with EC_50_s of 29.7 uM and 2.1 uM, respectively; Suppl. Table 2). There was no cytotoxicity observed against sub-confluent HFF (Suppl. Table 2), human fibroblast [25], or HepG2 cells (Suppl. Table 1).

### SAR studies of *C. parvum* inhibition and HCT-8 cytotoxicity

The isolation of a suite of related norditerpenes with a wide range of anti-parasitic and cytotoxic activities allowed us to identify some structural features correlated with biological activity. The tetracyclic norditerpene dilactones produced by fungi and plants are generally classified into three structural groups (Fig. 1). Type A norditerpenes contain an α-pyrone [8(14), 9(11)-dieneone], type B compounds include a 7α, 8α-epoxy-9(11)-dieneone and type C compounds have a 7(8), 9(11)-dieneone structure. Among the group of 14 *O. truncatum* compounds that we tested, there were examples in all three structural classes (albeit only a single “type A” example). We also want to emphasize the confusing nature of the trivial names of these compounds compared with their structure class (e.g. oidiolactones A and C are “type B” norditerpenes, while oidiolactone B is a “type A” norditerpene) along with references to rings A through D.

Comparison of the two most active compounds oidiolactones A (**1**) and B (**7**), shows that the presence of a 7,8 double bond vs. a 7α, 8α-epoxy group leads to a 2-fold increase in inhibitory activity against *C. parvum* and 2.6-fold increase against *T. gondii* (Fig. 2E). However, this structural difference also causes oidolactone B to be ∼15x more cytotoxic towards the host HCT-8 cells under non-confluent conditions (Suppl. Table 2). An α-OMe group at C-13 vs a hydroxy moiety also leads to higher levels of anti-parasitic activity (compare **1** to **3** and **7** to **10**). The presence of a hydroxy group on C-3 of the A ring eliminates activity for the epoxide derivatives (compare **1** to **4**) and significantly decreases parasite growth inhibition for the type C dieneones (compare **7** to **8**, and **10** to **9**). Opening of the δ-lactone ring (ring C) abolishes inhibitory activity for compound **5** with geminal O-methyl groups, and reduces activity for the only aldehyde derivative in the series, **6**. Opening of the □-lactone (ring D, compound **11**) strongly reduces activity. The only example of an α-pyrone norditerpene in this series, compound **12**, was inactive against the parasites (Fig. 1) and lacked cytotoxicity (Suppl. Table 2 and [25]).

Although oidiolactone B (**7**) was the most potent inhibitor of *C. parvum* in this series, oidiolactone A (**1**) had a higher therapeutic index when comparing the sub-confluent HCT-8 cytotoxicities (TI: EC_50_ *C. parvum*/EC_50_ HCT-8= 56) In further characterizing anti-parasitic activity in vitro and in vivo, we therefore focused on oidiolactone A as the most potent and potentially least cytotoxic compound.

### In vitro pharmacokinetics of oidiolactone A

There is no general consensus about whether oral cryptosporidium drugs should target intracellular (but extra-cytoplasmic) parasites in the gastrointestinal lumen directly or be absorbed systemically (ie. have high oral bioavailability). We assessed the in vitro stability of oidiolactone A with both human and mouse blood plasma and liver microsomes to identify potential liabilities and help predict its in vivo pharmacokinetics and develop a preliminary dosing regimen for in vivo studies. The compound was less stable in human vs mouse plasma, in contrast to most reports of small molecule in vitro stability comparisons across species[27], but more stable in human vs mouse liver microsomes (Table 1). We also determined that oidiolactone A exhibits high passive permeability using PAMPA (Pe 11.0 ± 1.3 × 10^−6^ cm/s), which is consistent with calculated lipophilicity and solubility values (Log P = 1.44 and Log S = 2.46). Oidiolactone A is not predicted to be a substrate or inhibitor of p-glycoprotein receptors.

**Table 1.** *In vitro* metabolic stabilities of oidiolactone A.

|  | Mouse plasma<br>stability t<sub>1/2</sub> (min) | Human plasma<br>stability t<sub>1/2</sub> (min) | Mouse microsomes<br>stability <sup>a</sup> t<sub>1/2</sub> (min) | Human microsomes<br>stability t<sub>1/2</sub> (min) |
| --- | --- | --- | --- | --- |
| oidiolactone A (**1**) | 108.0 ± 1.8 | 17.5 ± 0.1 | 110.4 ± 4.3 <sup>b</sup> | 317.7 ± 16.7 |
| verapamil | | | 2.8 ± 0.02 | 11.2 ± 0.2 |

Data are presented as mean ± SD. <sup>a</sup>CYP enzyme cofactor: NADPH. <sup>b</sup>Control in the absence of NADPH showed remaining percentage less than 80% at the end of incubation of 60min (Susceptible to non-NADPH dependent degradation).

### Oidiolactone A inhibits intracellular *T. gondii* replication, but does not have any apparent effect on parasite morphology

To determine if oidiolactone A had an effect on the morphology of replicating intracellular *T. gondii* parasites, treated parasites were evaluated by immunofluorescence assay. Intracellular parasites were treated for 24 hours with oidiolactone A or DMSO control, fixed and then visualized with anti-surface antigen 1 (SAG1) antibody. Treated parasites were morphologically similar to untreated parasites, but were present at much lower abundance (Fig. 3) as expected (see Fig. 2).

**Figure 3:**
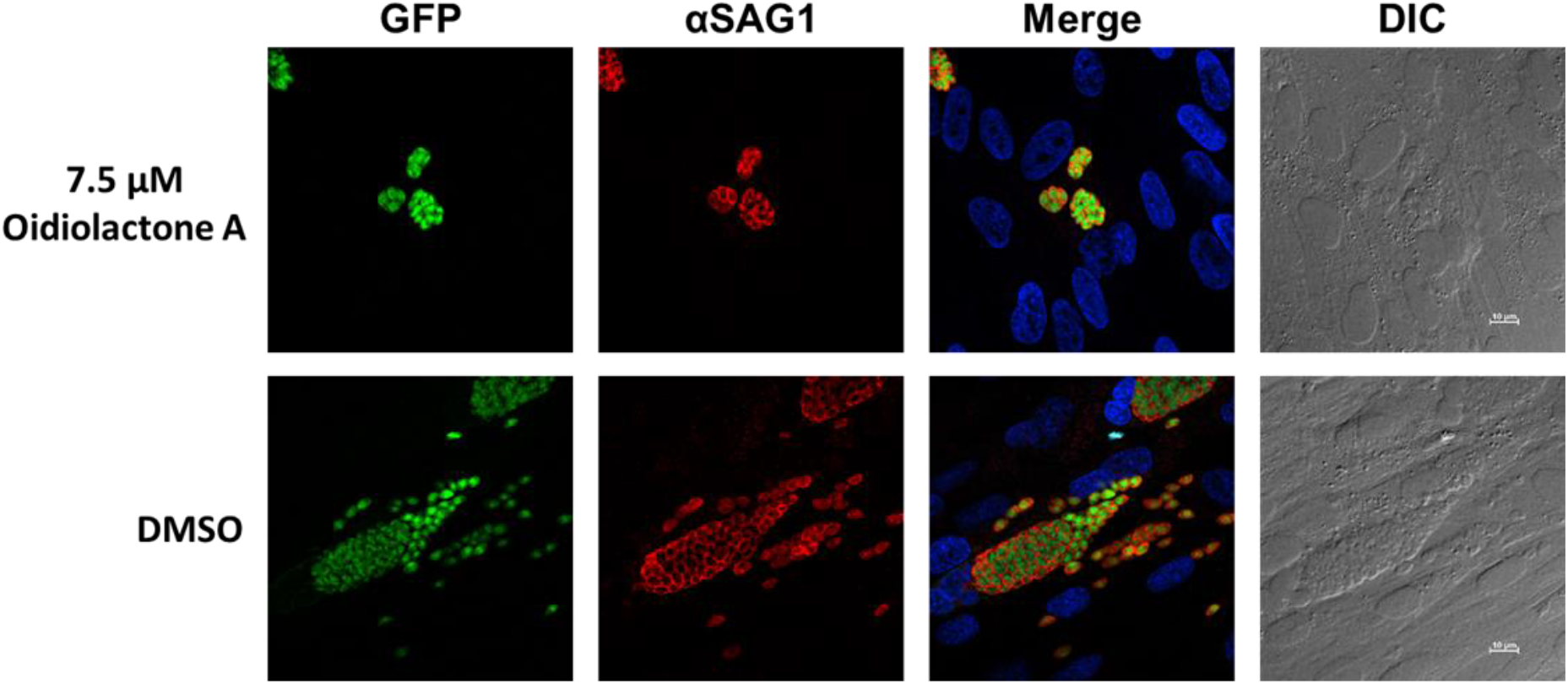
Oidiolactone A inhibits growth of *Toxoplasma gondii*. TgRH-Luc:GFP infected HFFs were treated with oidiolactone A for 24 hours. Cells were then fixed and labeled with anti-SAG1 antibody (red) to visualize the tachyzoite membrane and Hoescht (blue) to label host cell nuclei. Scale bar=10 µm.

### Oidiolactone A has -cidal activity against both *Cryptosporidium* and *Toxoplasma*

Since oidiolactone A inhibited parasite growth, but did not appear to overtly damage parasites (Fig. 3), intracellular Cp-NLuc and TgRH-Luc:GFP parasites were treated for varying lengths of time, from 4 to 24 hours, after which the compound was removed and the parasites were allowed to recover for a further 48 hours for Cp-NLuc and 24 hours for TgRH-Luc:GFP. After 8-12 hours of treatment with oidiolactone A at the EC_90_ (7.5 uM for Cp-NLuc and 15 uM for TgRH-Luc:GFP), neither Cp-NLuc and TgRH-Luc:GFP recovered, suggesting a -cidal versus -static effect (Fig. 4 A and B, respectively).

**Figure 4:**
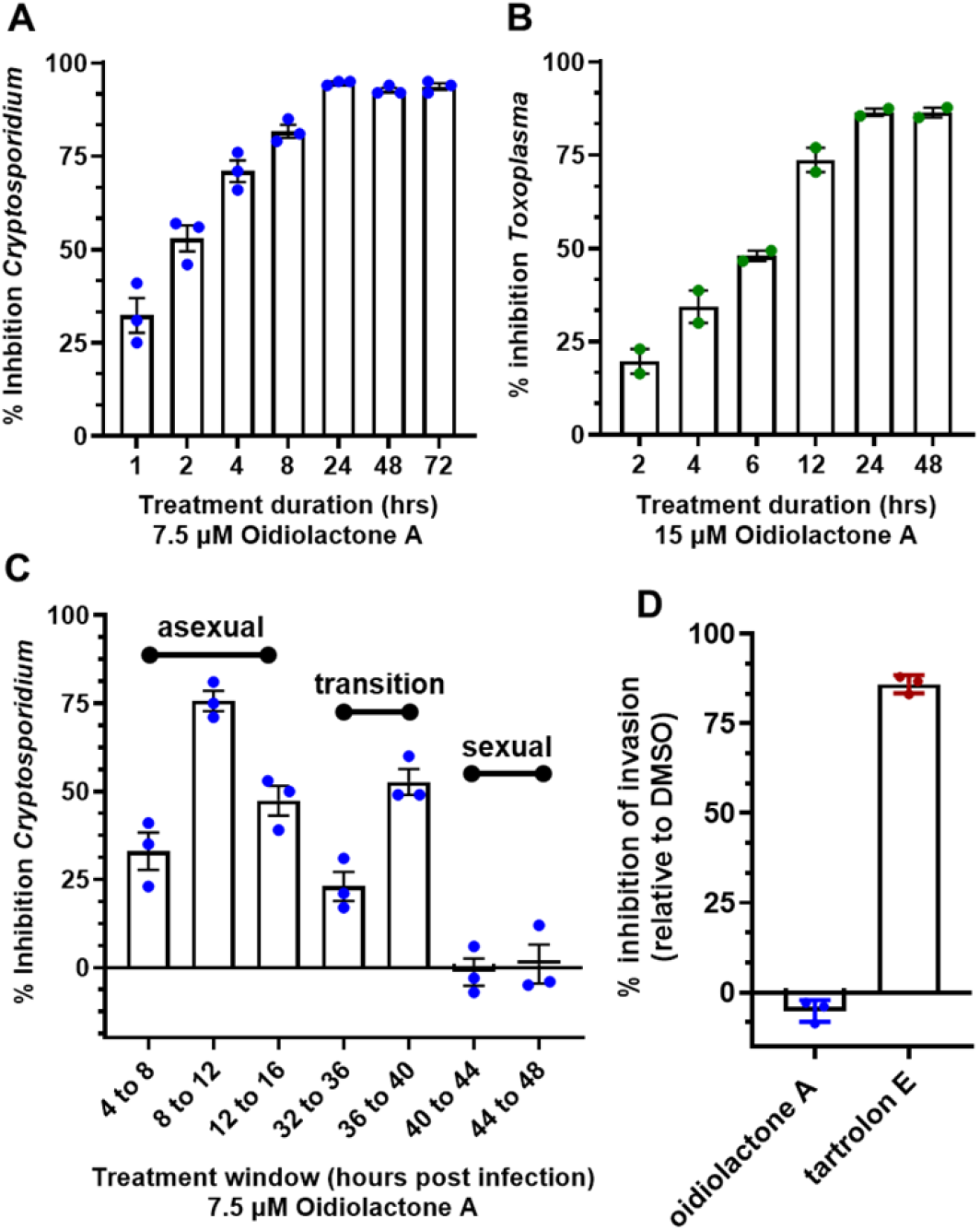
Oidiolactone A rapidly kills intracellular, asexual parasite stages, but does not inhibit sporozoite invasion into host cells. **(A)** Cp-NLuc was allowed to infect HCT-8 cells for 24 hours, at which point compound or DMSO vehicle control was added for the lengths of time indicated. At the end of each time point, compound was washed out and infection allowed to proceed until the plate was read at 72 hours post-infection. Results are the percent of growth inhibition relative to the DMSO control. **(B)** HFF cells infected with TgRH-Luc:GFP parasites were treated as described in (a) but the plate was read 48 hours post-infection. Results calculated as in A. **(C)** HCT-8 cells were infected with Cp-NLuc and the compound or DMSO added for 4 hour intervals as indicated. The plate was read at 72 hours post infection, and percent inhibition of parasite growth calculated based on DMSO control for that time interval. **(D)** Cp-NLuc oocysts were added to HCT-8 cells incubated in 7.5 µM oidiolactone A, 100 nM tartrolon E or DMSO and allowed to infect for 3 hours at which point plate was read for luciferase expression. Percent inhibition calculated by comparison to DMSO control. All data are compiled from 3 biological replicates. Error bars: +/-SD.

### Oidiolactone A inhibits *C. parvum* parasites during asexual replication and during the transition to sexual stages

While *T. gondii* tachyzoite cultures, used here to determine anti-*Toxoplasma* activity, are representative of just the one life cycle stage, *Cryptosporidium* in vitro cultures develop through three rounds of asexual replication during the first ∼32 hours, at which time they begin to transition to formation of the male and female gametes [4]. By 44 hours post infection the culture consists of primarily gametes. In standard 2D cultures, the life cycle halts at fertilization and no oocysts are produced. However, accurate timing of asexual and sexual development in HCT-8 cells has been reported [4], and this timeline can be used to evaluate the susceptibility of different parasite stages to anti-parasitic compounds [4,28]. Oidiolactone A (7.5 μM) was added to HCT-8 cells for 4 hour intervals post infection with NLuc-*C. parvum* oocysts. Inhibition (20-75%) was observed only for parasites in the asexual stage of proliferation, and during transition to sexual stages (Fig. 4C). Oidiolactone A exhibited no activity against gamete stages.

### Oidiolactone A does not inhibit excystation or prevent invasion of *Cryptosporidium* sporozoites into host cells

Compounds that can prevent excystation or sporozoite invasion into host cells could potentially be used as prophylactic agents, an application critical to prevent calf cryptosporidiosis and subsequent environmental contamination. Oidiolactone A had no effect on oocysts excystation (Suppl. Fig. 1A). The ability of oidiolactone A to inhibit invasion was evaluated by luciferase expression (Fig. 4D) and by immunofluorescence assay with enumeration of trophozoite stages (Suppl. Fig. 1B). Oidiolactone A did not exhibit any effect on parasite invasion in contrast to the control compound tartrolon E, which completely inhibited parasite invasion.

### Oidiolactone A reduces infection in IFN-γ knockout mice infected with Cp-NLuc

We evaluated the in vivo efficacy of oidiolactone A in a model of acute *C. parvum* infection. Five week old IFN-gamma knockout (GKO) mice were infected with mouse-adapted Cp-NLuc oocysts. On days 5 and 6 post-infection, mice were treated with four doses of 4 mg/kg of oidiolactone A by oral gavage, every 12 hours. Control mice were treated in parallel with the vehicle control. Fecal samples were collected daily, until day 14 post-infection, when the mice were euthanized, and serum was harvested for biochemical profiling. Over the course of infection, oidiolactone A reduced shedding of oocysts as determined by relative luminescence units (RLU) per gram of feces, (Fig. 5A) indicating that the treatment with oidiolactone A was effective in reducing the parasite load. The treatment resulted in an overall 70% reduction in *C. parvum* oocyst shedding (Fig. 5B), suggesting that this compound has therapeutic potential for treating cryptosporidiosis. Additionally, assessments of liver, renal, metabolic, and cardiac functions, as well as protein status and electrolyte levels, showed no statistically significant differences between the control and treated groups, indicating the absence of any signs of toxicity (Suppl. Table 3).

**Figure 5:**
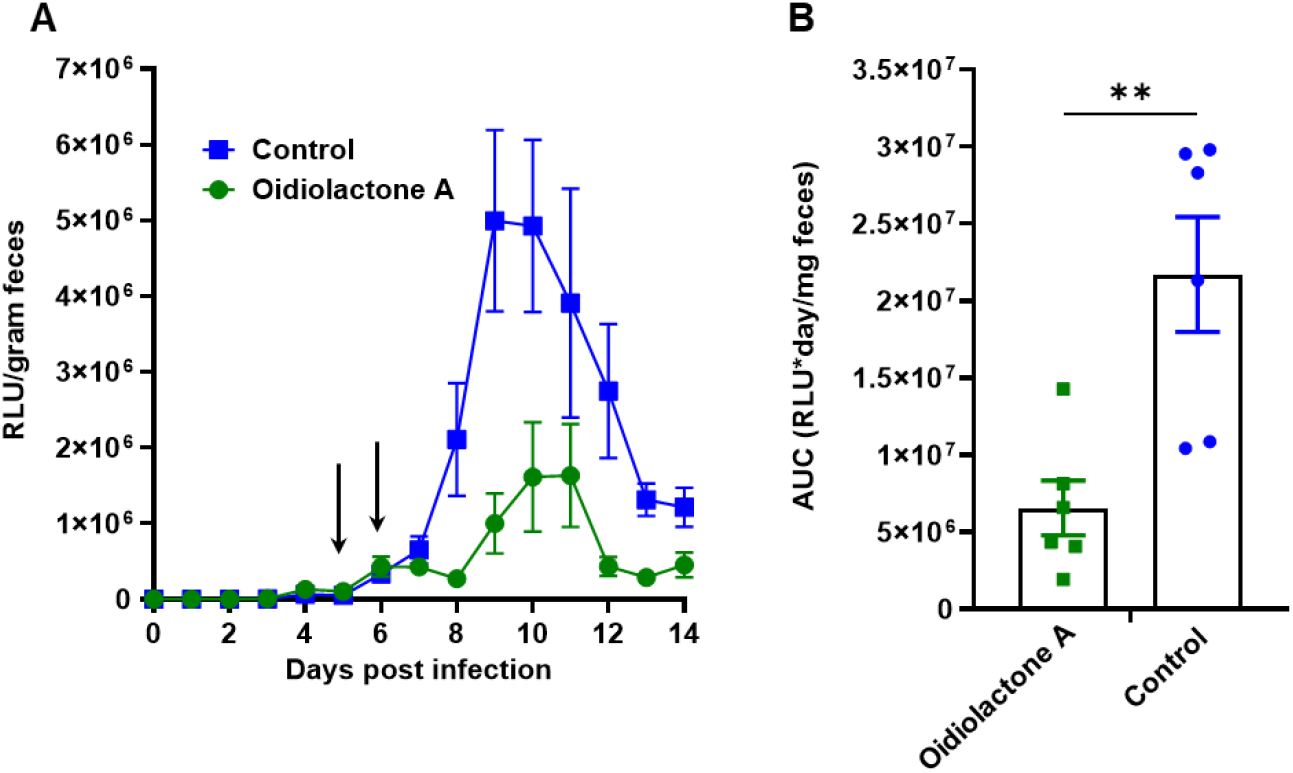
Oidiolactone A significantly reduces oocysts shedding in *Cryptosporidium* infected IFNγ-knockout mice. **(A)** Mice were infected with Cp-NLuc and oocyst shedding monitored daily by luciferase expression. On days 5 and 6 post infection, 4 mg/kg oidiolactone A was administered every 12 hours by oral gavage (arrows). Controls received DMSO in parallel. Error bars represent standard error of the mean. **(B)** The area under the curve (AUC) of the oocysts shedding for each mouse was calculated. Each point is data from one mouse. A Mann-Whitney t-test was used to compare the groups. **p=0.0087

## Discussion

Cryptosporidium remains one of the few, if perhaps only, globally distributed pathogens with no effective vaccines or therapeutics for prevention and treatment [29]. To date the number of compounds in the therapeutic pipeline for cryptosporidiosis are few, and the most promising target a single enzyme, raising the possibility that resistance could arise once the compounds are deployed in the clinic [14]. Several compounds are under advanced preclinical or clinical investigation [30–35, 60,61]. A repurposed leprosy drug, clofazimine, with anti-*Cryptosporidium* activity, failed in clinical trials due to solubility and bioavailability issues [36], highlighting the need for early pharmacokinetic analysis and multiple lead compounds. Very little investigation into natural products as a source of anti-*Cryptosporidium* compounds has been conducted, despite the fact that parasitic gregarines, close relatives of *Cryptosporidium*, are widely found in all environments [37] and may provide an ecological basis for the evolution of natural anti-parasitics [38]. Additionally, natural products have historically provided us with our most effective and robust anti-parasitic drugs [23,24]. Thus far, two purified natural compounds have been reported to have anti-*Cryptosporidium* activity. One of these, tartrolon E, isolated from a shipworm bacterial symbiont, has rapid killing activity against the parasite, is highly effective in vivo, and is also broadly effective against several apicomplexan parasites [38]. The marine sponge metabolite, leiodolide A, has potent submicromolar in vitro activity against *Cryptosporidium*, underscoring the potential of natural products as a source of therapeutics for cryptosporidiosis [39]. This report of norditerpene oidiolactones with anti-parasitic activity, identified in a very small phenotypic screen, further supports this approach.

The oidiolactone norditerpenes are members of a large, diverse group of tetracyclic dilactone terpenoid metabolites isolated from several fungal species and more than a dozen plant species in the coniferous Podocarpaceae family [40–42]. Some of the plant derived compounds have been studied for their potential ecological roles as insecticides, anti-feedants, herbicides and plant growth regulators [43–45]. Many of the fungal and plant norditerpenes have also been tested for potential medicinal applications, and have been reported to have broad anti-microbial activities (bacteria, fungal and parasitic) as well as anti-oncogenic and anti-inflammatory activity [42]. This study is the first report of members of this structural class having potent activity against the apicomplexan parasites *Cryptosporidium parvum* and *Toxoplasma gondii*. A suite of seven structurally related norditerpene lactones (=oidiolactones) with antiparasitic activity were identified from an initial in vitro growth inhibition screen. Both oidiolactones A and B had submicromolar EC_50_s against intracellular *Cryptosporidium* and *Toxoplasma*, with little cytotoxicity for mammalian cells. Since oidiolactone A exhibited less toxicity towards growing HCT-8 cells, further investigation of the anti-parasitic activity of the fungal norditerpenes was conducted with oidiolactone A.

Oidiolactone A killed replicating *Toxoplasma* and *Cryptosporidium* within 24 hours, and was primarily effective against asexual stages of *Cryptosporidium*. The compound did not inhibit sporozoite infection of host cells. Most critically, it was highly effective against the parasite in vivo in an immunocompromised mouse model of acute infection, even at a low dose (4 mg/kg). To evaluate the range of activity against other protozoan parasites, we also tested the compounds against the causative agent of malaria, *Plasmodium falciparum*, and the intestinal parasite *Giardia lamblia*. Oidiolactone A exhibited low micromolar activity against *P. falciparum* but did not inhibit *G. lamblia* (EC_50_ >30 μM).

There is no clear agreement on the ideal pharmacokinetic parameters for effective anti-cryptosporidial drugs. There is some data to suggest that optimal anti-cryptosporidial compounds would be retained in the gut with slow systemic uptake [46–48]. For example, the in vivo efficacy of a series of related bumped kinase inhibitor (BKI) pre-clinical lead compounds, ATP-competitive inhibitors of parasite calcium-dependent protein kinases, was found to depend solely on their concentration in the large intestine [48]. It is unclear if this would be true for all classes of anti-*Cryptosporidium* compounds. Retention in the intestine does minimize the need for plasma and microsomal stability and high bioavailability, and potentially reduces off target cytotoxicity effects [46]. The relatively short half life of oidiolactone A in mouse blood plasma (<2 hrs), high membrane permeability, and in vivo efficacy suggests that the compound may be acting through direct exposure in the intestine. An additional consideration for intestinal absorption of drugs is the involvement of P-glycoprotein (P-gp) located on the apical membrane of enterocytes in the intestinal epithelium[49]. These receptors function to pump substrates back into the intestinal lumen, effectively reducing their intracellular concentration and systemic absorption. Recent research with the BKIs found that derivatives that were not recognized by P-gp were more effective in vivo [50]. Oidiolactone A is not predicted to be a P-gp substrate or inhibitor[51,52], but additional testing using in vitro bidirectional permeability assays and co-administration studies with P-gp inhibitors will be needed for confirmation[50,53].

The discovery of a suite of 14 structurally related norditerpene lactone derivatives with widely different anti-parasitic activities provided an opportunity to compare the structure activity relationships (SAR) related to both anti-parasitic activity and host cell cytotoxicity (Fig. 6). For this series, the primary features of the most active compounds include a C-13 O-methyl ether, lack of a hydroxy group at C-3 and intact lactone rings. Although the type C diene congener, oidiolactone B, was the most potent inhibitor of *C. parvum* and *T. gondii*, it was also more toxic towards actively dividing host cells while the type B epoxide derivative, oidiolactone A, had a higher therapeutic index. This initial set of SAR data provides a useful set of guidelines for further testing and discovery of additional related natural norditerpene dilactones as well as synthetic derivatives.

**Figure 6:**
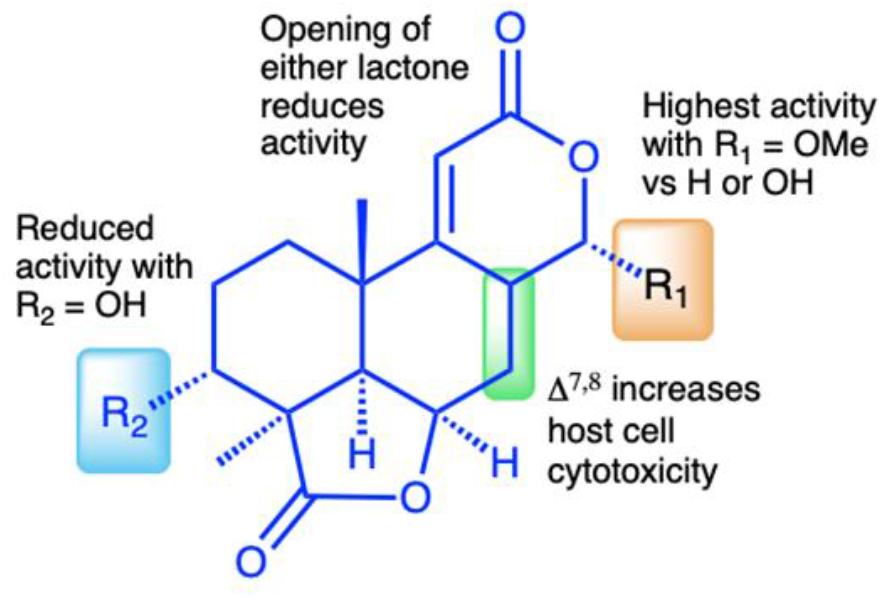
Structure activity relationship (SAR) summary for oidiolactones from *O. truncatum* tested against nluc *C. parvum* in HCT-8 cells. “Activity” refers to inhibition of parasite growth, and “cytotoxicity” refers to inhibition of host cell growth under sub-confluent conditions.

Oidiolactone A is also an attractive compound for optimization as it is produced in high concentrations by the fungus (∼70 mg/L) under unoptimized conditions, and has several functional group “handles” for semi-synthetic transformations. Because there has been significant interest in the potential anti-cancer activity of many norditerpene dilactones such as the nagilactones and podolactones, there is also a rich literature precedent for both semi-synthesis and de novo synthesis for several members of this class[54–56], providing established methods for production or modification. Additionally, several of the anti-cancer studies with related norditerpene dilactones included both oral and intraperitoneal administration of candidate compounds using mouse models, which provides some additional data about in vivo dosing, oral bioavailability, and toxicity[57–59].

Although the molecular target is not yet known, it is likely that the mechanism of action of the norditerpene dilactone metabolites is different than for other known cryptosporidial inhibitors because the structure is so distinct from reported anti-cryptosporidium compounds[30–36, 38,60,61]. Previously identified targets of oidiolactones and closely related analogs related to their anti-cancer activities include IL-1β and TNF-α[62], while the nagilactones and podolactones were found to induce apoptosis[58], inhibit protein synthesis[63] and inhibit transcription factor AP-1, activator protein 1[64,65]. Oidiolactone A inhibits intracellular replication of the parasites. It is as yet unclear if this activity is due to inhibition of a parasite-specific target or modulation of a host cell factor or process that the parasite requires for intracellular growth.

Oidiolactone A demonstrates strong anti-cryptosporidial activity both in vitro and in vivo and is readily accessible as a natural product and via well established synthetic methods. These data collectively provide compelling evidence of the potential of this class of norditerpenes as therapeutics for untreatable cryptosporidiosis.

## Materials and methods

### Compounds and extracts

The isolation and structural elucidation of compounds **1**-**9** and **11**-**14** were previously described[25]. An additional analog, yukonin (**10**), was also included in bioactivity screening. The fungus *Oidiodendron truncatum* (MN0802960) was isolated from the 25th level of the Soudan Iron Mine in Tower, Minnesota and cultured on potato dextrose agar for ten days. Approximately 1/8th of a fully colonized agar plate was chopped into small pieces and vortexed in 5 mLs of PBS in a 50mL conical tube and used to inoculate 200 g of rice media (50% rice/water by weight) in a 1L flask which was incubated for 30 days at room temperature (∼22°C). Methanol and ethyl acetate extracts (3 × 300 mL each) of the cultures were combined, dried, and successively partitioned with ethyl acetate, n-hexanes, and butanol. The ethyl acetate soluble fraction was separated using medium pressure flash chromatography (Teledyne ISCO Combiflash^®^ Rf; Solid phase Redisep^®^ Rf 24 g silica; gradient elution 0 – 100% of EtOAc/n-Hexane for 35 minutes; flow rate 25 mL/minute). Fractions were pooled into 18 fractions (**F1**-**F18**) based on TLC analysis. Fraction **F10** (260.4 mg) was further separated using semi-preparative HPLC (gradient elution 25 – 60% acetonitrile/H_2_O for 23 minutes; flow rate 3 mL/minute, column: Vision HT-C18 classic, 10 × 250 mm) to generate **F.10.11**. (12.2 mg). Fraction F.10.11 was then subjected to another HPLC separation (gradient elution 35 – 57.5% acetonitrile/H_2_O for 9 minutes; flow rate 1mL/minute, column: Altima HP-C18, 4.5 × 250 mm) to give yukonin (**10**, 1.25 mg) which was identified by comparison of NMR and mass spectrometry data to literature values[66].

### Parasites and host cells

The Iowa strain transgenic *Cryptosporidium parvum* oocysts expressing nanoluciferase (Cp-NLuc) were acquired from the University of Arizona, Tuscon, AZ (https://acbs.arizona.edu/cryptosporidium-production-laboratory). Oocysts were used within 3 months of acquisition. Before use, oocysts were bleached and washed as previously described [67]. The type I RH strain of *Toxoplasma gondii*, expressing green fluorescent protein and luciferase (TgRH-Luc:GFP) was obtained from Jeroen Saeij at the University of California, Davis. TgRH-Luc:GFP parasites were maintained in human foreskin fibroblasts (ATCC SCRC-1041TM) as described [68]. Human ileocecal colorectal adenocarcinoma cells (HCT-8, ATCC® CCL244™) were maintained as recommended and were subcultured no more than 30 times. *P. falciparum* cultures were maintained following a modified Trager and Jensen protocol[69]. The multi-drug-resistant *P. falciparum* line Dd2 was grown in RPMI 1640 supplemented with 25 mM HEPES pH 7.4, 26 mM NaHCO3, 2% dextrose, 15 mg/L hypoxanthine, 25 mg/L gentamicin, and 0.5% Albumax II in human A+ erythrocytes. Cultures were maintained at 4% hematocrit at 37 °C with 5% CO2.

### *Cryptosporidium parvum* growth inhibition assays

The in vitro growth inhibition assay of *C. parvum* has been previously described [39]. Briefly, clear-bottom, white-sided 96 well plates were seeded with HCT-8 cells. Once the cells reached between 50% and 75% confluency, they were infected with Cp-NLuc oocysts (10,000/well) in complete media containing 0.6% taurocholate. At 24 hours post infection (hpi) the infected cells were treated with compounds at various concentrations. DMSO was run in parallel as a vehicle control. Parasite growth was determined at 72 hpi by relative luminescence using Promega Nano-Glo substrate (Promega, Madison WI) and a SpectraMax® L Microplate Reader (Molecular Devices, San Jose, CA, USA). Percent inhibition of parasite growth was calculated as [(RLU_DMSO_ − RLU_compound_)/RLU_DMSO_] × 100. Each assay was run in triplicate with three biological replicates.

To determine -static versus -cidal activity, the infection assay was conducted as described, but infected cells were treated with compounds for 30 minutes up to 24 hours. The compounds were removed after the designated times, and infection quantified at 72 hours post-infection. To determine activity of the compounds against asexual and sexual stages of the parasite, compounds were added for four hour windows at times corresponding to asexual or sexual development. Parasite proliferation was then determined at 72 hours post-infection.

### *Toxoplasma gondii* growth inhibition assays

The *T. gondii* inhibition assay has been described previously [38]. Briefly, HFF cells were seeded onto a white-sided, clear-bottom 96 well plates, grown to confluence and 5,000 TgRH-Luc:GFP parasites added to each well. At 24 hpi, compounds or DMSO were added. The growth of the parasites was determined by relative fluorescence at 48 hpi using Promega Bright-Glo (Promega, Madison WI) substrate. Plates were read and percent inhibition of parasite growth calculated as described above.

Determination of -static versus -cidal activity was conducted as indicated above, with parasite growth quantified after 48 hours of infection. Each assay was run in triplicate with three biological replicates.

### *Plasmodium falciparum* growth inhibition assay

Antiplasmodial EC_50_ results were determined using a fluorescence-based SYBR Green I assay performed using asynchronous cultures[70]. For screening, parasites were diluted to 1% parasitemia and 2% hematocrit, then incubated with serial dilutions of compounds in microtiter plates for 72 h at standard growth conditions. Plates were subsequently frozen at −80 °C to facilitate lysis. After thawing, plates were incubated with 1× SYBR Green I in lysis buffer (20 mM Tris-HCl, 0.08% saponin, 5 mM EDTA, 0.8% Triton X-100) for 45 min at room temperature. Fluorescence was measured at an excitation wavelength of 485 nm and emission wavelength of 530 nm in a Synergy Neo2 multimode reader (BioTek, Winsooki, VT, USA). Relative fluorescence units (RFUs) were normalized based on chloroquine-treated and no treatment controls. Serial dilutions of compounds were prepared in RPMI with final assay conditions of ≤0.2% DMSO.

### HCT-8 and HFF cytotoxicity assays

Cytotoxicity of the compounds was determined against both confluent (the conditions of the infection assays) and growing HCT-8 and HFF host cells as described[25, 38]. Cells were seeded into white-sided, clear-bottom 96-well plates, and were grown to either 40% or 100% confluency. Compounds were added to cells at various concentrations up to 100 uM in a final DMSO concentration of 0.5%. DMSO was run in parallel as a vehicle control. After 48 hours, the viability of the cells was determined by either quantification of cellular ATP (Cell-Titer Glo cell viability assay; Promega, Madison WI)[38] or by MTT assay[25]. Viability was determined 24 hours after compound addition to HFF cells or 48 hours after compound addition to HCT-8 cells or HepG2 cells. The percent cell growth was then calculated with the following formula: % Cell Growth = [RLU_compound_/RLU_DMSO_] × 100.

### HepG2 Cytotoxicity Assay

Mammalian cell cytotoxicity was assessed in HepG2 human hepatocytes using an MTS 3-(4,5-dimethylthiazol-2-yl)-5-(3-carboxymethoxyphenyl)-2-(4-sulfophenyl)-2*H*-tetrazolium)-based cytotoxicity assay. For cytotoxicity testing, ∼2250 cells were seeded into each well of a 384-well microtiter plate, and the cells were incubated for 24 h. Serial dilutions of the compounds were added in triplicate starting at 25 μM (final concentration). Cells were further incubated for an additional 48 h at 37 °C in an atmosphere containing 5% CO2.

Zero percent growth control wells were incubated with 5% Triton X-100 for 5 min prior to MTS addition. Following the addition of MTS to all wells, the plates were incubated for an additional 4 h under the same environmental conditions before taking absorbance measurements. Absorbance values were recorded at 490 nm using a Synergy Neo2 multimode reader (BioTek), and values were normalized based on Triton X-100-lysed and no treatment controls.

### Compound inhibition assay against *Giardia lamblia* WBC6 trophozoites

Growth inhibition assays were performed using wildtype *G. lamblia* WBC6 cells at 37°C under hypoxic conditions in a tube base susceptibility assay described below. *G. lamblia* WBC6 WT trophozoites were grown in TYI-S-33 medium supplemented with 10% bovine serum and 0.05mg/mL bovine bile[71]. A 3-fold serial concentration of pure fungal inhibitors starting at 29 µM was screened against 75000 cells/ml of parasites in modified TYI-S-33 medium with final dimethyl sulfoxide (DMSO) concentration of 0.14%. Metronidazole was run in parallel at a final highest concentration of 66.7 µM and 3-fold subsequent serial dilutions. Untreated negative control contained 0.14% of DMSO. Sealable test tubes with a total volume of 700 µL were used to perform this assay. The test tubes were incubated at 37°C for 72-hours. After incubation, the tubes were iced and then the reaction was transferred to 96-well plates. In the plate, 50 µL of an ATP binding luminescence marker, CellTiter-Glo (Promega, Madison, WI) was added to each well and allowed to develop at room temperature for 10 minutes. The assay was read and analyzed using an EnVision Plate Reader (Perkin Elmer, Waltham, MA). Response curves and EC_50_ values (the concentration at which 50% of *G. lamblia* trophozoite growth is inhibited) were calculated using GraphPad Prism (GraphPad, LaJolla, CA).

### *Cryptosporidium* excystation and invasion assays

Oocyst excystation rates in the presence of compounds was determined as previously described[67]. The *C. parvum* invasion inhibition assays were conducted in two formats. In the first format, described previously [72], HCT-8 cells were seeded on glass slides (CELLTREAT Scientific Product, Pepperell, MA, USA) and allowed to grow to confluence. Oidiolactone A was added to the cells for one hour. TrtE and DMSO were run in parallel as positive and negative controls. The cell monolayers were then infected with 1.2×10^6^ wild type *C. parvum* oocysts. After 3 hrs of incubation at 37°C, the slides were washed five times with 111mM D-galactose, fixed with 4% paraformaldehyde and permeabilized with 0.25% Triton-X. *C. parvum* sporozoites were detected with rabbit anti-gp40 [73] and Alexa Fluor 594 goat anti-rabbit antibodies. The HCT-8 cell nuclei were stained with 0.09mM Hoechst. Slides were coded to enable unbiased counting, and sporozoites from at least 10 randomly selected fields were counted in each sample at 100x magnification (Zeiss^®^ Axioscope 5). The percentage of invaded parasites was determined using the following equation: % Invaded Parasites = [parasites_compound_/parasites_DMSO_] × 100. This experiment was run in duplicate with 2 biological replicates.

In the second format, HCT-8 cells were seeded onto clear bottom, white sided 96 well plates and grown to confluence. Compounds or DMSO were added to the cells for 1hr. The cell monolayers were then infected with 2×10^5^ Cp-NLuc oocysts and incubated and washed as described above. Relative luminescence was determined using the Promega Nano-Glo kit and the SpectraMax^®^ L Microplate Reader. The percentage of invaded parasites was determined using the following equation: % Invaded Parasites = [RLU_compound_/RLU_DMSO_] × 100. The experiment was run in triplicate with three biological replicates.

### Inhibition of *Cryptosporidium* infection in IFN-gamma knock-out (GKO) mice

Twelve female C57BL/6 IFN KO mice (4 weeks old) from Jackson Laboratory were housed in a controlled environment with a 12-hour light/dark cycle and access to food and water ad libitum. Cp-NLuc oocysts were isolated from infected mouse feces and purified using a sodium chloride density gradient [26]. Mice were inoculated via oral gavage with a suspension of 10^4^ oocysts in 50 µL sterile PBS. On days 5 and 6 post-infection, mice were treated with 4 mg/kg of oidiolactone A or DMSO vehicle control every 12 hours for a total of four doses (6 mice per treatment group). Fecal samples were collected daily and stored at −20 °C until analysis. Feces were homogenized, centrifuged, and the supernatant was incubated with Nano-Glo® substrate in a 96-well plate as described [26]. Luminescence was measured in a microplate reader immediately after substrate addition. Oocysts shedding based on RLUs/gram of feces were plotted over time and the area under the curve determined for each mouse.

Mice were sacrificed 14 days post-infection, and serum was collected for biochemical analysis. The biochemical analyses were performed using the Beckman Coulter AU480 automated analyzer (Beckman Coulter, Brea, CA) at the Clinical Pathology Laboratory, Veterinary Medical Center, University of Minnesota.

### Statistics

The inhibition curves and EC50s were determined using the log[inhibitor] vs. response-variable slope (four parameter) regression equation of GraphPad Prism v10.0.1. Differences between groups were determined by parametric or non-parametric t-tests as indicated in figure legends.

### Pharmacokinetics

#### Plasma stability assay

The plasma stability assay was performed in duplicate by incubating each selected compound (1 µmol/L final concentration) in normal pooled human plasma (Innovative Research, Novi, MI, USA) at 37 °C. At 0, 1, 3, 6, and 24 h, a 40 µL aliquot of the plasma mixture was taken and quenched with 120 µL of acetonitrile containing 0.1% formic acid. The samples were then vortexed and centrifuged at 15,000 rpm (Thermo Scientific Sorvall ST 8R, NI, Germany) for 5 min. The supernatants were collected and analyzed by LC-MS/MS to determine the *in vitro* plasma half-life (*t*_1/2_).

#### Microsomal stability assay

The *in vitro* microsomal stability assay was conducted in duplicate in commercially available human liver microsomes (Sekisui XenoTech, Kansas City, KS, USA), which were supplemented with nicotinamide adenine dinucleotide phosphate (NADPH) as a cofactor. Briefly, a compound (1 µmol/L final concentration) was spiked into the reaction mixture containing liver microsomal protein (0.5 mg/mL final concentration) and MgCl_2_ (1 mmol/L final concentration) in 0.1 mol/L potassium phosphate buffer (pH 7.4). The reaction was initiated by addition of 1 mmol/L NADPH, followed by incubation at 37 °C. A negative control was performed in parallel without NADPH to reveal any chemical instability or non-NADPH dependent enzymatic degradation for each compound. A reaction with positive control verapamil was also performed as in-house quality control to confirm the proper functionality of the incubation systems. At various time points (0, 5, 15, 30 and 60 min), a 40 µL of reaction aliquot was taken and quenched with 120 µL of acetonitrile containing 0.1% formic acid. The samples were then vortexed and centrifuged at 15,000 rpm for 5 min at 4 °C. The supernatants were collected and analyzed by LC-MS/MS to determine the *in vitro* metabolic half-life (*t*_1/2_).

#### PAMPA membrane permeability assay

The membrane permeability of selected compounds was evaluated using the Corning^®^ BioCoat™ Pre-coated PAMPA Plate System (catalog. No. 353015, Corning, Glendale, AZ, USA). The pre-coated plate assembly, which was stored at −20°C, was taken to thaw for 30 min at room temperature. The permeability assay was carried out in accordance with the manufacturer’s protocol. Briefly, the 96-well filter plate, pre-coated with lipids, was used as the permeation acceptor and a matching 96 well receiver plate was used as the permeation donor. Compound solutions were prepared by diluting the 10 mmol/L DMSO stock solutions with DPBS to a final concentration of 100 μmol/L. The compound solutions were added to the wells (300 μL/well) of the receiver plate and DPBS was added to the wells (200 μL/well) of the pre-coated filter plate. The filter plate was then coupled with the receiver plate and the plate assembly was incubated at 25 °C without agitation for 5 h. At the end of the incubation, the plate was separated and the final concentrations of compounds in both donor wells and acceptor wells were analyzed using LC-MS/MS. Permeability of the compounds were calculated using the Eq. (1):

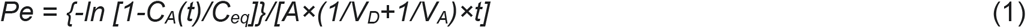

where A = filter area (0.3 cm^2^), V_D_ = donor well volume (0.3 mL), V_A_ = acceptor well volume (0.2 mL), *t* = incubation time (seconds), C_A_(*t*) = compound concentration in acceptor well at time *t*, C_D_(*t*) = compound concentration in donor well at time *t*, and C_eq_ was calculated using the Eq. (2):

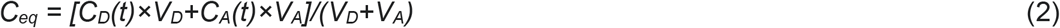

A cutoff criteria of *Pe* value at 1.5 × 10^−6^ cm/s was used to classify the compounds into high and low permeability according to the literature report of this PAMPA plate system[74].

The logP and logS values were calculated in silico using the SwissADME server[75], and analysis of P-gp substrate/inhibitor likelihood was determined using the PgpRules prediction server[51].

## Declarations

**All authors have seen and approved the manuscript. This manuscript has not been accepted or published elsewhere**

## Ethics Statement

All animal studies described in this manuscript were approved by the University of Minnesota IACUC (protocol #2110-39522A). Animals were euthanized by isoflurane inhalation followed by cervical dislocation as is consistent with the current AVMA Guidelines for the Euthanasia of Animals.

