## Supplemental Data for "Norditerpene natural products from subterranean fungi with anti-parasitic activity"

**Supplemental Table 1: Activity of oidiolactones against *Plasmodium falciparum* isolates and HepG2 cells**

|  | **EC_50_** | | **TC_50_** |  |
| --- | --- | --- | --- | --- |
| **Compound** | **Dd2 (μM)** | **3D7(μM)** | **HepG2 (μM)** | **SI** |
| 1 oidiolactone A | 2.263 ± 0.460 | 3.327 ± 0.640 | > 25 | >11 |
| 2 oidiolactone C | 2.489 ± 0.198 | 2.777 ± 0.397 | 17.056 ± 0.821 | 7 |
| 3 oidiolactone D | > 5 | > 5 | > 25 | >5 |
| 4 oidiolactone G | > 5 | > 5 | > 25 | >5 |
| 5 oidiolactone H | n.t. | n.t. | n.t. |  |
| 6 oidiolactone I | > 4.985 | 3.651 ± 0.596 | > 25 | >5 |
| 7 oidiolactone B | 0.983 ± 0.111 | 1.217 ± 0.419 | 10.334 ± 2.185 | 11 |
| 8 oidiolactone J | > 5 | > 5 | > 25 | >5 |
| 9 oidiolactone K | > 5 | > 5 | > 25 | >5 |
| 10 yukonin | n.t. | n.t. | n.t. |  |
| 11 oidiodendronic acid | > 5 | > 5 | > 25 | >5 |
| 12 oidiolactone L | > 5 | > 5 | > 25 | >5 |
| 13 oidiolactone E | > 5 | > 5 | > 25 | >5 |
| 14 LL-Z1271β | > 5 | > 5 | > 25 | >5 |

Data are presented as the mean +/- SD. SI: selectivity index=EC_50_ HepG2/ EC_50_ compound

n.t.: not tested

**Supplemental Table 2: Cytotoxicity of oidiolactones against confluent and subconfluent host cells**

| Compound | HCT-8  (confluent) | HCT-8  (sub-confluent) | HFF  (confluent) | HaCAT (sub-confluent)^a^ |
| --- | --- | --- | --- | --- |
| 1 oidiolactone A | >30 | 29.7 | >30 | >30 |
| 2 oidiolactone C | >30 | >30 | >30 | >30 |
| 3 oidiolactone D |  |  |  | >30 |
| 4 oidiolactone G |  |  |  | >30 |
| 5 oidiolactone H |  |  |  | >30 |
| 6 oidiolactone I | >30 | 4.8 | >30 | >30 |
| 7 oidiolactone B | >30 | 2.1 | >30 | 21.4 |
| 8 oidiolactone J | >30 | >30 | >30 | >30 |
| 9 oidiolactone K |  |  |  | >30 |
| 10 yukonin | >30 | 2.5 | >30 | n.t. |
| 11 oidiodendronic acid | >30 | 19.7 | >30 | >30 |
| 12 oidiolactone L |  |  |  | >30 |
| 13 oidiolactone E |  |  |  | n.t. |
| 14 LL-Z1271β |  |  |  | >30 |

TC50 values are in µM; n.t.=not tested. ^a^TC50 values for *Homo sapiens* HaCAT fibroblast cell line were previously reported in Rusman et al 2020 [25].

**Supplemental Table 3: Biochemical analysis of serum from infected and treated IFNγ -/- mice**

|  | **Sample ID** | | | |
| --- | --- | --- | --- | --- |
|  | **Oidiolactone A mouse 1** | **Oidiolactone A mouse 2** | **Control mouse 1** | **Control mouse 2** |
| BUN (mg/dL) | 25 | 18 | 19 | 20 |
| Creat (mg/dL) | <0.2 | <0.2 | <0.2 | <0.2 |
| Ca (mg/dL) | 10.6 | 10.5 | 10.9 | 10.5 |
| Phos (mg/dL) | 10.0 | 10.5 | 11.7 | 10.3 |
| Mg (mg/dL) | 2.8 | 2.7 | 3.2 | 3.0 |
| TP (g/dL) | 5.0 | 4.7 | 5.1 | 5.0 |
| Alb (g/dL) | 3.4 | 3.2 | 3.4 | 3.4 |
| Glob (g/dL, calc.) | 1.6 | 1.5 | 1.7 | 1.6 |
| Na (mmol/L) | 149 | 150 | 152 | 151 |
| Cl (mmol/L) | 112 | 113 | 112 | 112 |
| K (mmol/L) | 5.2 | 5.1 | 5.3 | 5.9 |
| HCO3 (mmol/L) | 15.9 | 13.8 | 17.1 | 16.8 |
| Osmol (calc.) | 311 | 309 | 317 | 313 |
| An Gap (calc.) | 26 | 28 | 28 | 28 |
| T. Bili (mg/dL) | 0.1 | 0.1 | 0.1 | 0.2 |
| ALP (U/L) | 171 | 193 | 161 | 182 |
| GGT (U/L) | <3 | <3 | <3 | <3 |
| ALT (U/L) | 42 | 33 | 22 | 36 |
| AST (U/L) | 207 | 252 | 113 | 211 |
| CK (U/L) | 1884 | 1352 | 817 | 1297 |
| Gluc (mg/dL) | 291 | 269 | 338 | 295 |
| Chol (mg/dL) | 98 | 87 | 114 | 98 |
| Amy (U/L) | 513 | 399 | 627 | 537 |
| LIH | 0 0 2 | 0 0 1 | 0 0 0 | 0 0 0 |

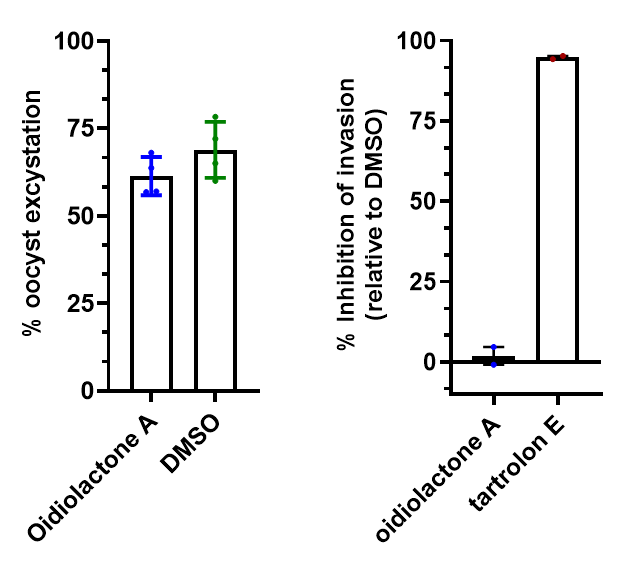

**Supplemental Figure 1: Oidiolactone A has no effect on oocyst excystation or sporozoites invasion into host cells. A.** Oocysts were excysted in buffer containing 7.5uM odiolactone A or DMSO vehicle control. After a 1 hour incubation, the % of oocysts that had excysted was determined. Data is compiled from 4 biological replicates (n.s.=not significant) **B.** WT *C. parvum* oocysts were added to HCT-8 cells in buffer containing 7.5 µM oidiolactone A, 100 nM tartrolon E or DMSO and allowed to infect for 3 hours at which point cells were washed, fixed and stained with anti-gp40 and Hoechst. Slides were coded for unbiased counting of invaded parasites. Percent inhibition calculated by comparison to DMSO control. Data represents 2 biological replicates.
